# Comparing the representation of a simple visual stimulus across the cerebellar network

**DOI:** 10.1101/2022.09.12.507660

**Authors:** Ot Prat, Luigi Petrucco, Vilim Štih, Ruben Portugues

## Abstract

The cerebellum is a highly conserved structure of the vertebrate central nervous system that plays a role in the timing and calibration of motor sequences. Its function is supported by the convergence of fibers from granule cells (GCs) and inferior olive neurons (IONs) onto Purkinje cells (PCs). Theories of cerebellar function postulate that IONs convey error signals to PCs that, paired with the contextual information provided by GCs, can be used as a teaching signal to guide motor learning.

Here, we use the larval zebrafish to investigate (i) how sensory representations of the same stimulus vary across GCs and IONs and (ii) how PC activity reflects these two different input streams. We use population calcium imaging to measure the cell responses to flashes of diverse luminance and duration to show that IONs and GCs encode different stimulus properties. First, most GCs show tonic and graded responses, as opposed to IONs, whose activity peaks only at on and off luminance transitions, in agreement with the notion that GCs and IONs encode context and error information, respectively. Secondly, we show that GC activity is patterned over time: some neurons had sustained responses for the entire duration of the stimulus, while in others activity was ramping up with slow time constants. This suggests that, by performing temporal integration, GCs could provide a basis that PCs may use to decode time. Finally, we show how PC activity can be largely reconstructed by a linear combination of granule cells and inferior olive neurons. Together, our observations give support to the notion of an error signal coming from IONs, and provide the first experimental evidence for a temporal patterning of GC activity over many seconds.

## Introduction

In order to orchestrate appropriate motor reactions and maximize survival chances of the organism, brains need to generate efficient sensory representations of the environment and its changes. The cerebellum has long been considered one of the main regions involved in the integration of sensory and motor representations, and plays a central role in motor coordination, the control of fine motor skills, and the calibration of reflexes and motor learning (Ito, 2006). These abilities are supported by the convergence of two information streams onto Purkinje cells (PCs): parallel fibers and climbing fibers (Eccles et al., 1967).

Parallel fibers originate in the granule cells (GCs) of the molecular layer of the cerebellum, the most abundant cell type in the human brain (Williams and Herrup, 1988). Thousands of parallel fibers establish excitatory synapses onto a single PC. Inspired by these numbers, the first theories of cerebellar function proposed that GCs sparsely code and expand the dimensionality of inputs, which originate in pre-cerebellar nuclei and arrive via mossy fibers (Marr, 1969; Albus, 1971). This high-dimensional representation of sensory information allows selective and differential weighting of stimulus properties and the acquisition of context-dependent cerebellar-guided behavioral responses (Dean et al., 2010). The wide scope of modalities that can be represented by GCs encompasses sensorimotor (Knogler et al., 2017), somatosensory (Arenz et al., 2009) and even predictive (Giovannucci et al., 2017; Wagner et al., 2017) information, but recent studies have challenged the notion of sparse representations (Giovannucci et al., 2017; Knogler et al., 2017; Wagner et al., 2017). In addition to sparse coding, theories postulate that GCs can provide temporal information to PCs, as feedforward control of movement depends on the ability of the cerebellum to perform temporally-specific learning (Kawato and Gomi, 1992; Ohyama et al., 2003) although see (Johansson et al., 2014). The large number of GCs makes them suited to encode this information, but while temporal patterning of GC responses has been assumed in multiple theoretical studies (Buonomano and Mauk, 1994; Medina and Mauk, 2000; Medina et al., 2000) it has found little empirical evidence so far, (but see (Kennedy et al., 2014)).

The second stream arriving to PCs via climbing fibers originates in inferior olive neurons (IONs). Each PC is innervated by a single climbing fiber (Eccles et al., 1966), that fires at low frequency and evokes large and sustained depolarizations. Most widespread theories argue that climbing fibers convey error signals (Ito, 2013). These, paired with the contextual information provided by parallel fibers, can guide motor learning (Marr, 1969; Albus, 1971) through LTD at the parallel fiber to Purkinje cell synapse (Ito, 2000).

To characterize and compare the functional properties of these convergent information streams, we studied them in the olivo-cerebellar circuit of the zebrafish larvae. The zebrafish cerebellum shares the same basic circuitry with the mammalian cerebellar cortex (Bae et al., 2009; Hamling and Tobias, 2015), but contains a greatly reduced number of neurons, and its cells are already mature by 5 days post-fertilization (dpf) (Hibi and Shimizu, 2012). Moreover, the fish cerebellum has been implicated in several sensory-motor behaviors such as the optokinetic reflex (Portugues et al., 2014) and optomotor response adaptation (Ahrens et al., 2012; Markov et al., 2021).

Taking advantage of the optical transparency of the fish larvae and the availability of transgenic lines that express in specific cell types of the circuit, here we sequentially monitored whole-field luminance responses of all main cerebellar subpopulations in awake and sensing zebrafish larvae. These experiments allowed us to study how GCs and IONs convey different information about the same stimulus. We observed that while GCs showed sustained responses carrying accurate information about the current state of the sensory input, IONs responded mostly to stimulus changes. Moreover, we were able to observe the signs of temporal integration in GCs responses, suggesting that their activity could provide a basis for temporal coding in the cerebellum. Finally, we investigated how GC and ION response profiles are integrated at the level of PCs.

## Results

### Experiment description and anatomy

We exploited the optical transparency and the genetic amenability of the larval zebrafish to follow how information over a simple visual stimulus is processed and transformed through different elements of the cerebellar circuitry. We employed two-photon microscopy in restrained zebrafish larvae to monitor neuronal activity while presenting them with visual stimuli projected on a screen below (Figure 1A). Larvae expressed GCaMP6s or a modified slow version of GCaMP6f (GCaMP6fe05) in either GCs, PCs or IONs (see Methods). The entire volume of either the cerebellum or the inferior olive was scanned in 1 μm (for inferior olive) or 1.5 μm (for the cerebellum) - spaced planes. In this way, we were able to acquire the responses of a large fraction of the GCs, PCs or IONs populations (Figure 1B).

**Figure 1:**
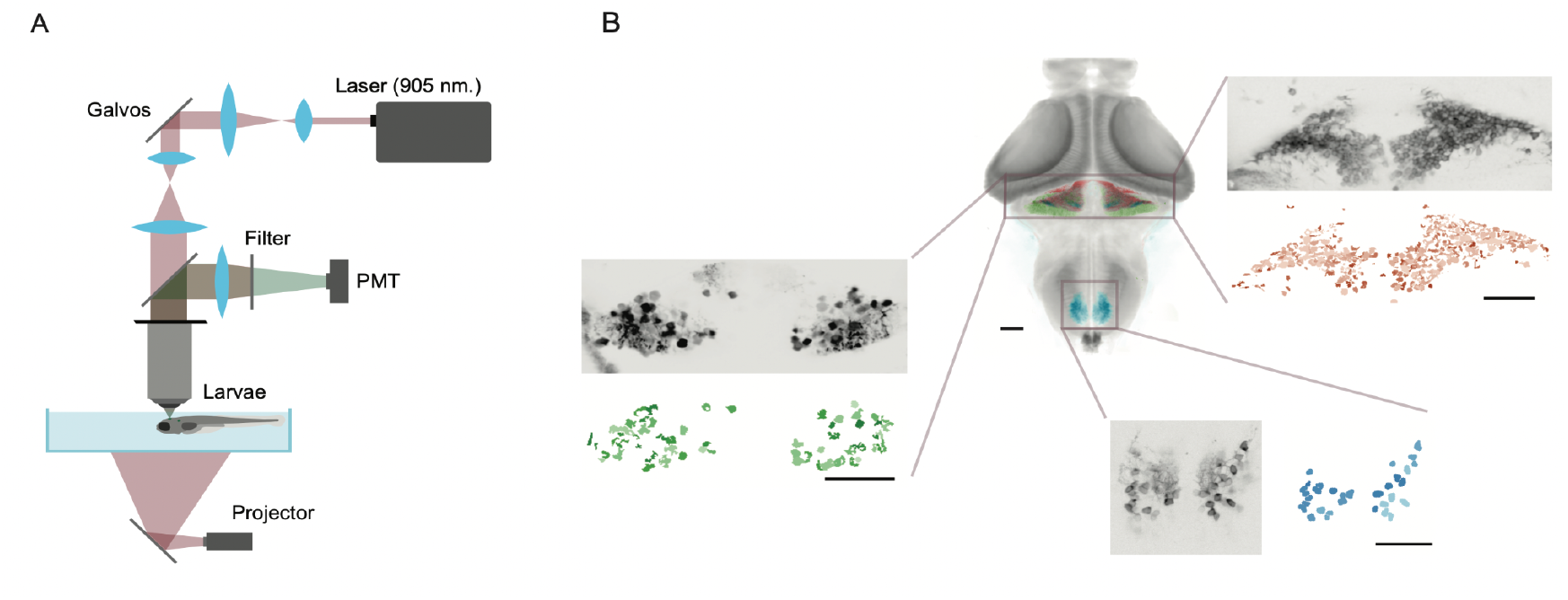
Experiment description and anatomy. A) Experimental setup: a two photon microscope was used to image 7 dpf head-restrained larvae while flashes of different luminance were projected on a screen below. B) Composite image showing stacks from GCs (red), IONs (in blue) and PCs (green) registered on a reference anatomy with the whole brain of a 7 dpf larva (in gray). In the enlargements, examples of the raw anatomy (in gray) and of the segmented ROIs from a 2-photon imaging session (in red, blue and green for GCs, IONs and PCs, respectively).

Zebrafish larvae are highly visual animals, and previous work from our laboratory (Knogler et al., 2017) has shown that luminance is very salient for neurons in the fish cerebellum. Therefore, we decided to investigate the diversity of responses across cell types as whole-field luminance was changed over a screen placed below the fish. This protocol elicited only a very mild behavior (Figure S1A), slightly reducing bout probability after light-to-dark transitions (Figure S1B-D), allowing us to analyze the sensory representations of the stimulus without the confounding effect of motor-related activity.

### Responses of GCs and IONs to different luminance levels

In a first set of experiments, we investigated the responses of GCs and IONs during a protocol where luminance was changed between four distinct levels. The progression was designed to ensure that transitions between all pairs of luminance levels were sampled (Figure S2A). The entire sequence was presented six times during the imaging of every plane, yielding a robust number of repetitions for all ROIs.

#### Granule cells

We collected the responses of 5013 ROIs from 5 larvae. To estimate how many cells were reliably engaged by the stimulus, we calculated the average correlation of calcium activity during each pair of stimulus trials (“reliability score”). In the GC population, the distribution of the obtained reliability scores was clearly bimodal (Figure 2A) with a fraction of about half strongly responsive cells (47.2% with an automatic thresholding, see Methods).

**Figure 2:**
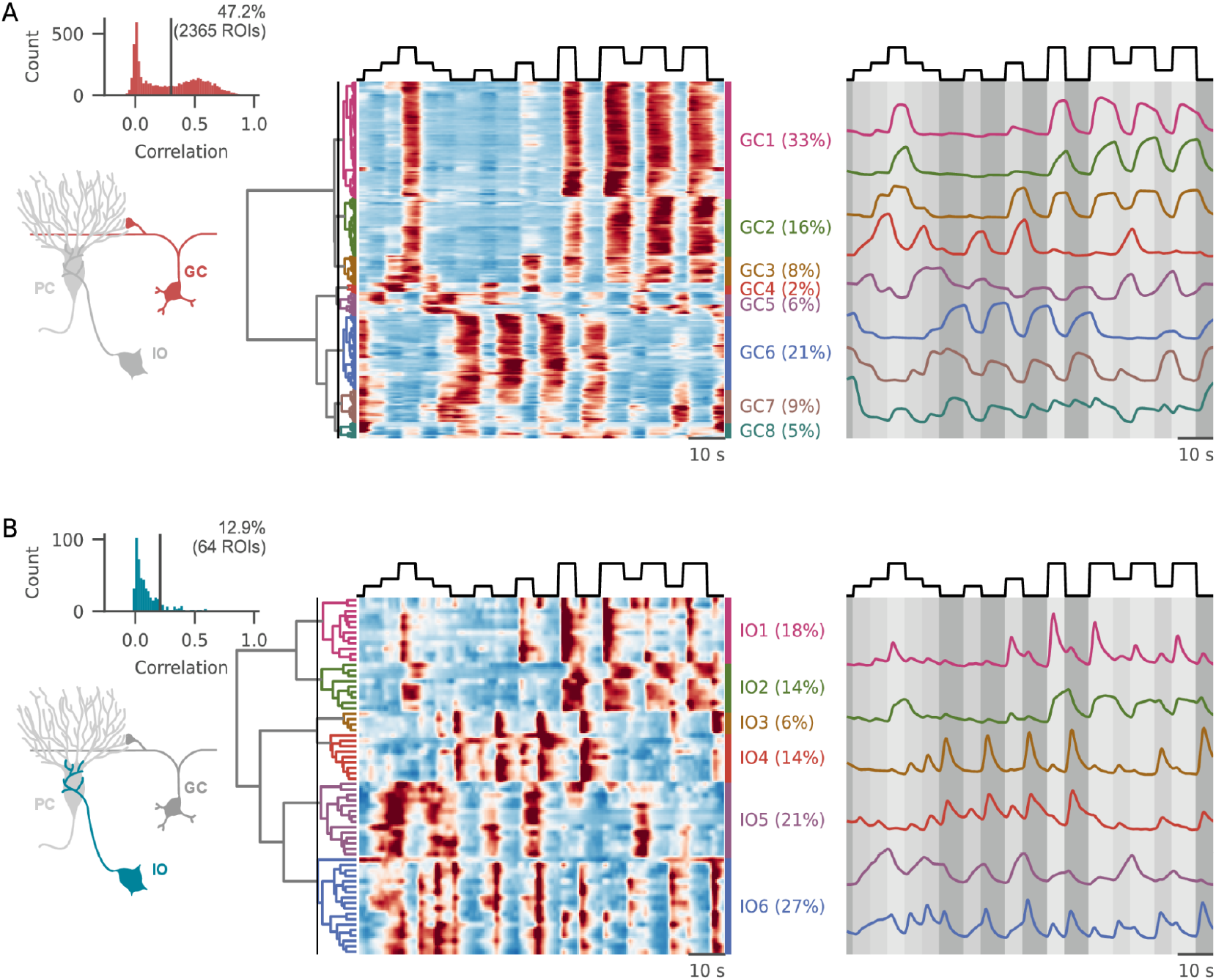
Responses of GCs, IONs and PCs to the “luminance steps” protocol. A) (*Left*) Histogram of the average inter-trial correlation (average correlation between activity recorded in different trials). The calculated threshold and the relative fraction of active cells are reported on the histogram. (*Center*) Average traces grouped after hierarchical clustering. Cutting the dendrogram at the height marked by the line (on the left) resulted in eight different clusters, whose average activity is shown on the right, over imposed on a shade matching at each timepoint the brightness level displayed. B) Same plot as in A), for IONs.

To explore the diversity of GC responses to the stimulus, we employed hierarchical clustering to classify the average response of all luminance-selective neurons (Figure 2A). The main branching in the distance tree arose from the distinction between ON and OFF cells, hereby defined as the property of an ROI of exhibiting a positive or negative calcium transient in response to an increase in the luminance level. The ON and OFF populations represented 57% and 43% of the responsive ROIs, confirming previous reports from our lab (Knogler et al., 2017). However, the presentation of intermediate luminance steps revealed the presence of a minority of ROIs (clusters GC4-5) that were specifically recruited by intermediate levels of luminance while silent at either minimum or maximum luminance (Figure 2A, Figure S2B).

ON and OFF cells did not respond in a homogeneous way to the battery of stimuli. First of all, we note that granule cell activation was almost always sustained during the entire presentation of each luminance step. Secondly, cells in both the ON and OFF clusters largely differed in terms of their threshold and saturation points (Fig S2B). Finally, for some clusters the activation during the presentation of an intermediate luminance level strongly changed depending on the previous luminance level (Fig S2C). We did not find any consistent anatomical organization in the localization of the clusters across fish.

#### Inferior olive

We then turned to the analysis of the responses of IONs (495 cells from 5 fish). IONs were less selective for the luminance stimulus, with a smaller fraction of cells (12.9% with automatic thresholding) exhibiting reliable responses across stimulus presentations. Even if the number of luminance selective cells was smaller, we could observe a broad diversity of responses in the IONs, with several types of response profiles appearing across fish (Figure 2B). In striking opposition with the sustained responses observed in GCs, ION responses were generally transient, peaking immediately after luminance transitions and quickly decaying afterwards. Some IONs were selective to either OFF or ON-transitions (clusters ION1 and ION3), while others were recruited by both (ION4). Only a minority of cells showed a sustained activation during high luminance periods (ION2, 14%), while two other clusters showed sustained activation during intermediate levels of luminance (ION5 and ION6). Interestingly, in cluster ION6, neurons seemed to combine a sustained activation during intermediate luminance levels with a marked transient activation upon stimulus transitions involving luminance decrements.

Taken together, these observations suggest that the two cerebellar input streams might convey complementary information to PCs. On one hand, GCs almost always presented sustained activation during the luminance steps, with a broad diversity of responses coming from different thresholds and saturation points. On the other hand, most IONs reacted to sharp transitions in the presented stimulus.

### GC and IONs respond to stimulus intensity and derivative, respectively

To further investigate this point, we carried out a regression-based analysis on the individual cell responses to better describe these different behaviors. We created a panel of regressors of two different kinds. The first group included regressors obtained by applying different gamma corrections to the raw luminance profile to describe responses with different thresholds and saturation points, plus a regressor obtained by subtracting two luminance traces with different gamma corrections to account for intermediate-luminance selective cells. The second group included regressors created from the luminance derivative: on-transitions, off-transitions, or both (Figure 3A). We then looked at the distribution of the best predictor for each GC and ION in the dataset. While GCs show large correlations with luminance levels and small correlations with transition-related regressors, IONs were divided in a group showing high luminance related correlations and a group showing high transition-related correlations. Overall, a larger fraction of IONs responses were better predicted by transition-related regressors compared to GCs responses (42.4% of IONs vs. 3% of GCs, Figure 3B-C). If our observations are true, it should be possible to accurately predict absolute luminance values (the only instantaneous property of the stimulus) from GC activity, and more accurately predict transitions of the stimulus from ION activity. By using a nonlinear decoder (a radial-basis-function support vector machine regressor - SVM), we tried to decode the current luminance level from the activity of IONs, or from subsets of GCs matching in size the ION population. This confirmed that the displayed luminance can be decoded to a higher degree of accuracy from GCs compared to the IO population, which has more transitory and inconsistent responses (Figure 3D).

**Figure 3:**
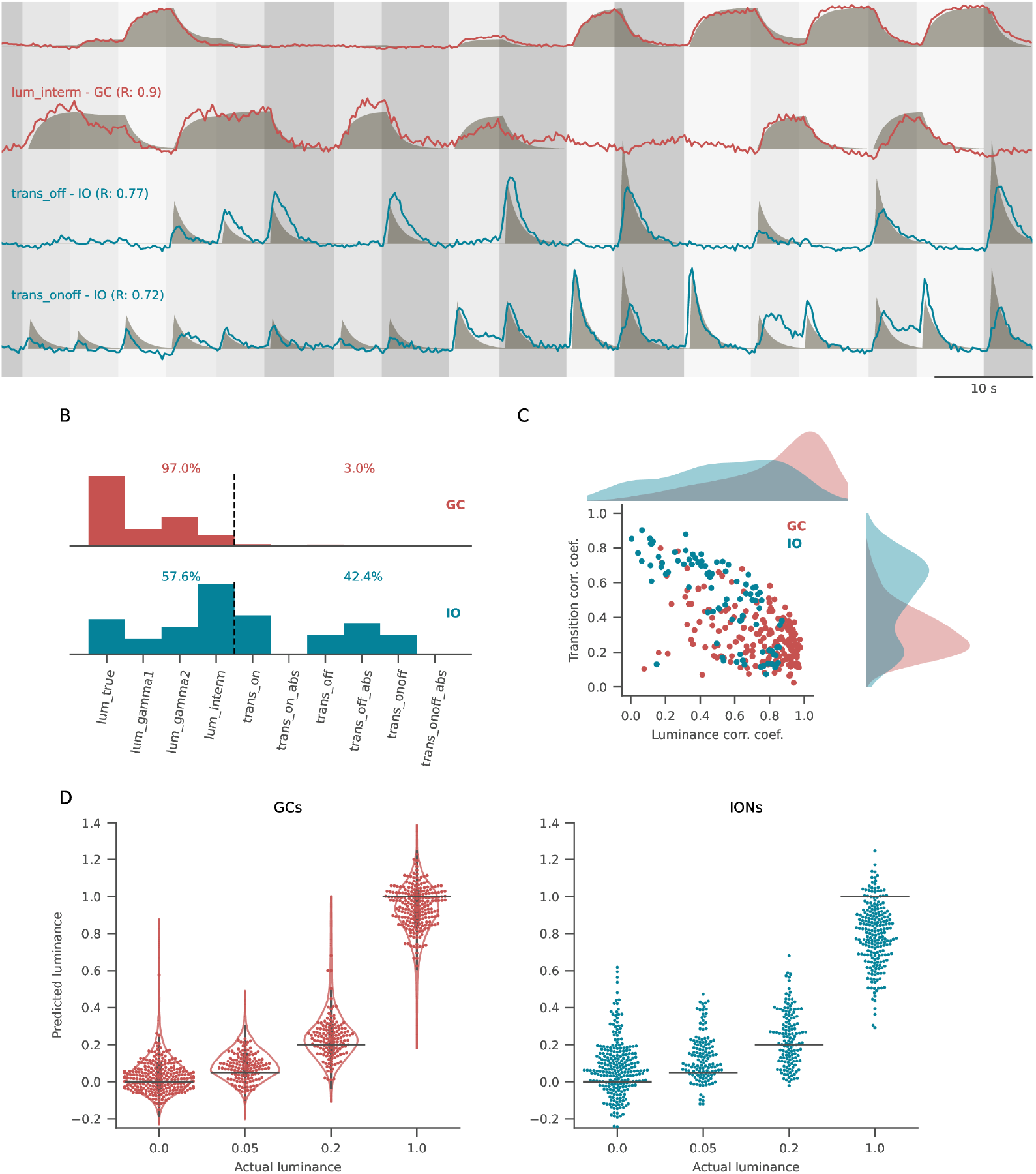
GC and IONs respond to stimulus intensity and derivative. A) Regressor-based analysis was used to calculate correlation coefficient between each cell fluorescence (lines) and a panel of regressors (shades). The plot shows the cells with the highest correlation values for luminance-related (first two rows) and transition-related (second two rows) regressors. B) Histogram of the distribution of best fitting regressors for GCs (top) and IONs (bottom). Regressors on the left of the dashed line are luminance related, regressors on the right are transition related. GCs score higher in luminance-related regressors compared to IONs. C) Scatterplot of the best transition-related coefficient and the best luminance-related coefficient for GCs (red) and IONs (blue), and their relative marginal distributions. The GC population has been downsampled randomly to match the number of IONs, while the marginal distributions refer to the entire population. GCs cluster in the bottom right quadrant of the plot (high luminance-regressors correlation, low transition-regressors correlation), while IONs show higher correlation coefficients for transition-related regressors. D) Performance of a non-linear decoder used to predict luminance values from GCs (left) and IONs (right) activity. Each point in the swarm plot is one frame of the protocol with the corresponding luminance level (horizontal line). For GCs, 20 iterations of the decoding analysis were performed, with the number of GCs downsampled to match that of IONs. The violin plots in the left panel show the average distribution of predictions across all iterations, while the dots correspond to a single representative iteration.

### Responses of GCs and IONs to flashes of different duration

Looking at the temporal dynamics of the GC responses, we noticed that in some clusters fluorescence was still markedly increasing at the end of the luminance step (cluster GC2, Figure 2A). We reasoned that this could be the hallmark of temporal integration of the luminance stimulus. Temporal integration has been suggested as a timing mechanism that the cerebellum could exploit to keep track of the period elapsed since stimulus onset, as is required for the acquisition of appropriately timed cerebellar-dependent responses such as eye blink delay conditioning (Medina and Mauk, 2000). Long time constants in GC responses have been postulated in models to account for timing in the cerebellum (Bullock et al., 1994), but they have never been convincingly observed experimentally.

To unravel temporal integration in the responses of the cerebellar circuitry, we designed a second protocol, consisting of three luminance flashes of different durations (3, 7 and 21 seconds) at maximal luminance (Figure S4A). The complete protocol was presented 6 times in each plane, and the reliability of responses of individual GCs and IONs was assessed as described above.

#### Granule cells

We sampled 13763 GCs from 5 fish and selected a fraction of 26% responsive ROIs for successive analyses (Figure 4A). As in the previous protocol, the major split in the GC diversity dendrogram corresponded mapped the difference between ON and OFF cells, (54% and 46% of the responsive cells, Figure 4A). Interestingly, the population of ON cells contained three clusters whose profiles differed in their temporal dynamics. Cluster GC3 showed the simple sustained activity that was reported in the steps protocol; cluster GC2 rapidly peaked after the flash onset, and then decayed to baseline before stimulus end; and cluster GC4 exhibited the response profile expected from a temporal integrator, ramping up slowly for the whole duration of the stimulus presentation. Since the experimental paradigm featured only sustained high luminance periods, we could not compare the kinetics of the luminance ON and the luminance OFF clusters.

**Figure 4:**
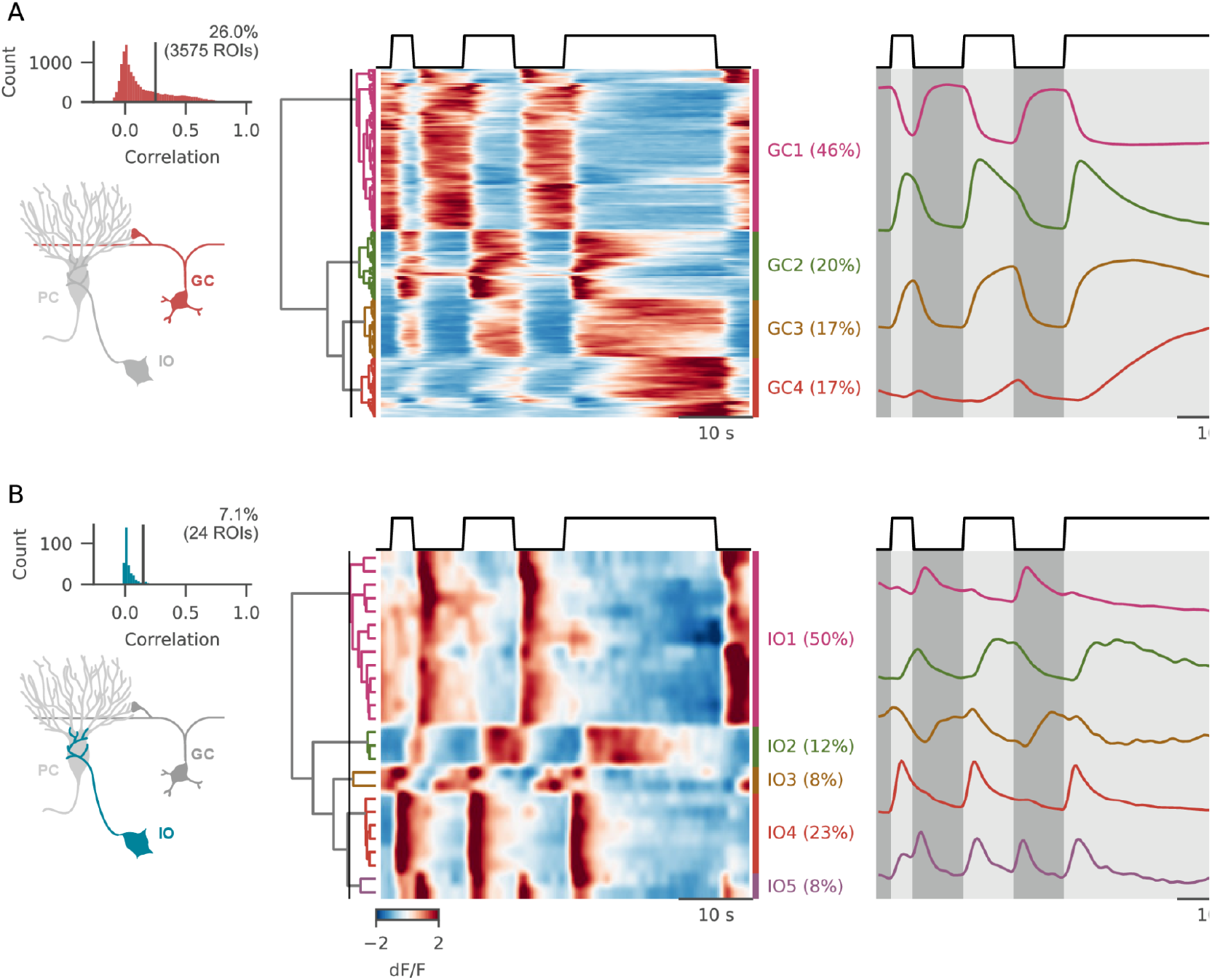
responses of GCs, IONs and PCs to the “flashes” protocol. A) (*Upper-left*) Inter-trial correlation histogram of GCs for the “flashes” protocol. (*Center*) Cells sorted after hierarchical clustering and (right) average activity for each cluster. B) Same plot as in A), for IONs.

#### Inferior olive

We imaged 337 IONs from 5 fish and, consistent with the previous observations, reliably responsive cells (7.1%) were mostly activated by luminance transitions. 50% were selective for negative luminance transitions (cluster ION1), 23% were selective for positive transitions (cluster ION4) and 8% for both transition types (cluster ON5). Only 20% of cells had sustained ON or OFF activations (clusters ION2 and ION3 respectively).

### Granule cells activity is temporally patterned

Intrigued by the slow temporal dynamics in cluster GC4 (Figure 4A), we decided to further investigate their temporal response properties. Different GCs responded with very different profiles that were consistent across stimulus presentations (Figure 5A). Remarkably, some cells from cluster GC4 clearly showed a period of suppressed activity after the stimulus presentation, followed by a positive, “integrator-like” response that could start rising several seconds after the beginning of the stimulus (Fig 5A, third trace). This might reflect the interplay between excitatory and inhibitory drives having different time constants, whose summing effects give rise to the observed diversity in the GCs responses timing.

**Figure 5:**
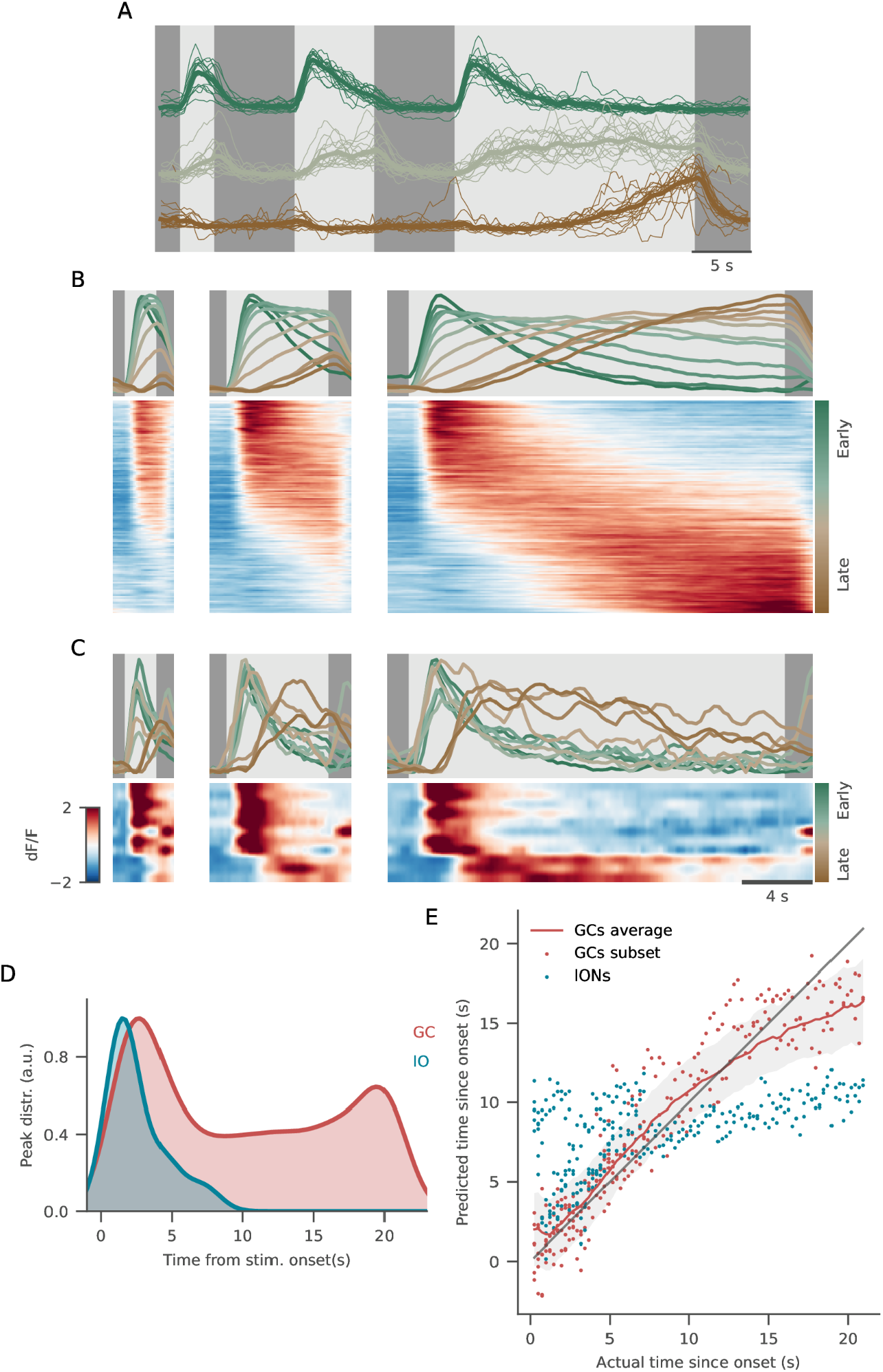
Granule cell activity is temporally patterned. A) Single traces for early-(green), sustained-(gray) and late-responding (brown) ROIs. The average response (thick line) is superimposed to all single repetitions (thin lines) of stimulus presentation. B) GCs included in any of the luminance-excited clusters were then sorted based on their center of mass (COM) during the longest flash, with earlier-responding neurons on the top, and late responding ones in the bottom (bottom panels). The above traces represent the average response for ROIs binned according to the time at which their COM was reached (green for early-responding neurons, brown for late-responding ones). C) Same plot as in B), for IONs. In this case, traces in the top panels correspond to single ROI responses. D) Probability density function describing the time points at which responses of neurons from the two imaged populations reach their maximal response. E) Scatter plot of predicted vs. actual time elapsed stimulus onset. Prediction was performed at each frame from IONs activity (blue dots) and from activity of a random subset of GCs (red dots). The red line and shaded area show average +/-standard deviation of predictions from 200 GCs subsets.

To look at the distribution of time constants in the responses of GCs from ON clusters (GC2, GC3, and GC4), we sorted them by their cross-validated center of mass (COM) computed on the 21-second flash (see Methods and Figure S5A). We observed a continuum between the different response profiles, ranging from early responding neurons that peak after the flash onset to late-responding neurons with responses slowly ramping up during stimulus presentation (Figure 5B). Most of the neurons reached their maximal activity shortly after stimulus onset, while many others were still ramping up at the time of its offset (Figure 5D). IONs did not show any integrating-like responses, and most ON cells peaked immediately after luminance onsets (Figure 5C-D).

Temporal integration would provide PCs with a representation of time elapsed since stimulus onset, as postulated in models of temporal learning in the cerebellum (Medina and Mauk, 2000; Yamazaki and Tanaka, 2009). Therefore, we examined whether it is possible to linearly decode this time since from the activity of an equal number of GCs or IONs. We observed that this decoding is possible to a high degree of accuracy by using the activity of granule cells (Figure 5E), with an R^2^ value of around 0.8. Since the temporal dynamics are nonlinear and share a saturating trend at longer durations, the optimal linear decoder overestimates the time at shorter durations and underestimates longer durations (Figure 5E). The inferior olive cells on the other hand show no long-term temporal patterning and it was therefore not possible to decode the duration of the stimulus presentation (R^2^=0,29, Figure S5B).

We conclude that the GCs exhibit diverse temporal dynamics that could provide PCs with a temporal basis for reading out the time since stimulus onset.

### PC responses to luminance stimuli

Both parallel fibers from GCs and climbing fibers from IONs make synaptic contact with PCs. Therefore, we decided to investigate the responses of PCs to the same battery of stimuli, to understand how the afferent cerebellar inputs are integrated when they converge at the level of their postsynaptic target. We used a transgenic line that expresses GCaMP6s selectively in all PCs. It is important to note that the unique biophysics of the PC spiking properties (namely, the difference between complex spikes elicited by climbing fibers and the simple spiking modulated by parallel fibers) might complicate the mapping between intracellular calcium-related fluorescence and spiking activity. Nevertheless, previous data from our lab (Knogler et al., 2019) has shown that both complex spikes and bursts of simple spikes induce comparable GCaMP6s fluorescence signals in fish PCs.

We first analyzed the responses of PCs to the “steps” protocol (Figure 6A) in 3318 ROIs from 5 fish. Of these, 20.3% were recruited by the stimulus. Given the well characterized nature of synaptic inputs of PCs, we expected to find response profiles reflecting the profiles observed in the GC and the ION populations. PCs showed both ON- and OFF-selectivity, a fraction of which were selective for intermediate luminance levels, as was observed in GCs (cluster PC1). However, there were significantly more

**Figure 6:**
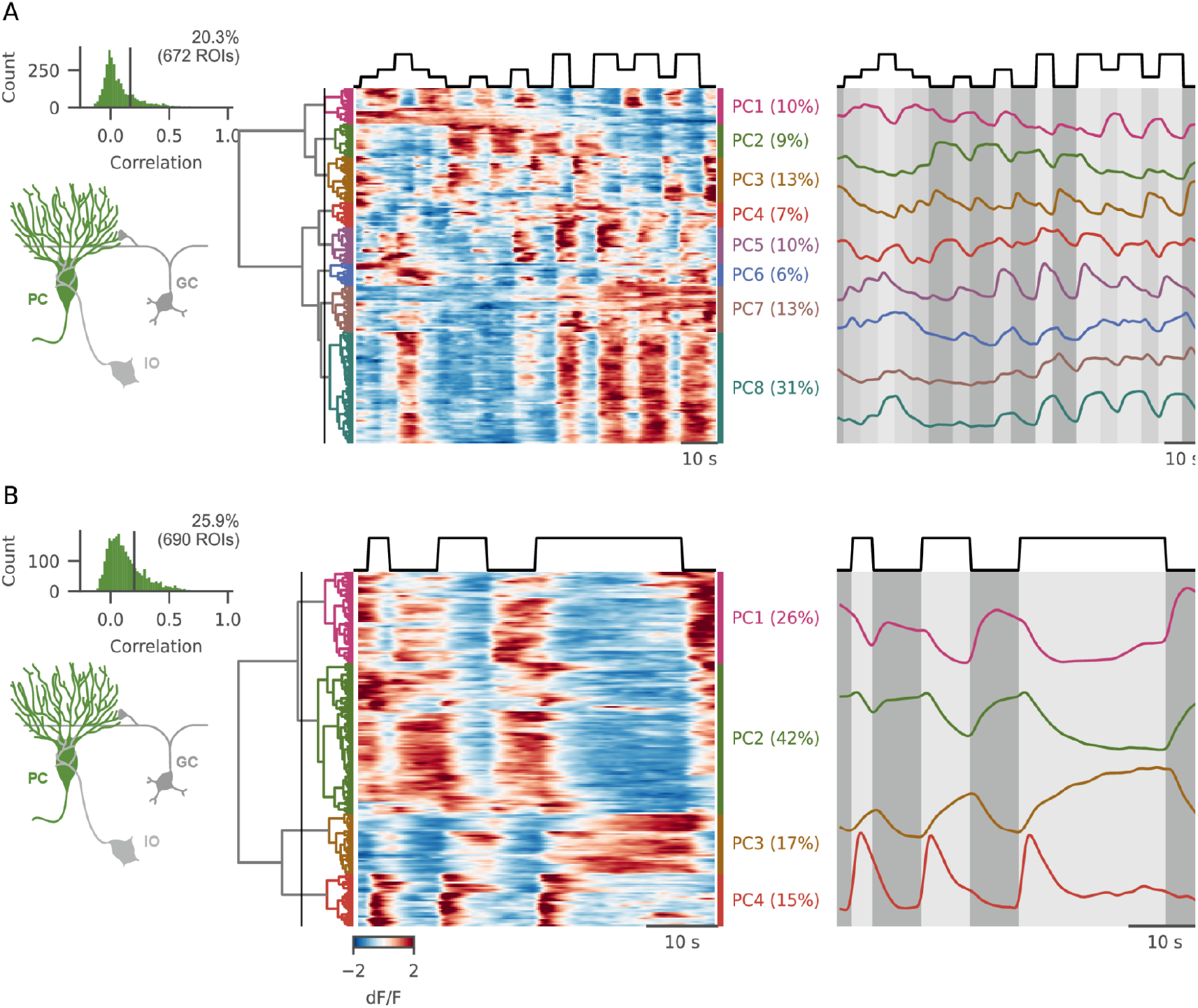
PC responses to luminance stimuli. A) (*Upper-left*) Histogram of the average inter-trial correlation and its relative threshold (*center*) Hierarchical clustering of PC responses to the “steps” protocol, and average activity of the clusters selected after cutting the dendrogram at the height marked by the lines. Shade matches at each timepoint the brightness level displayed. B) Same as in A), for the “flash” protocol.

ON cells than OFF cells (67% vs 32%, respectively) suggesting that the former are over-represented in PCs compared to GCs. Importantly, from the clustering approach we could find only one definite group of cells showing transient, transition-related activity (PC3), while most clusters looked modulated mainly by GC inputs.

Then, we turned to the investigation of PC activity during the “flashes” protocol (Figure 6B), analyzing the responses of 2267 ROIs from 5 fish, (25.9% responsive). PC activation profiles showed mixed IO and GC features. Just as in GCs, there were many late-responding ROIs that exhibited integrating-like responses (cluster PC3), but no cluster showed a constant, sustained response throughout the stimulation period.

Moreover, a large group of ON cells showed only a transient ION-like activation after luminance onset (cluster PC4) that was faster than the one observed in GCs (cluster GC2 in Figure 4), suggesting an input from IONs only.

### Modeling of PC responses from GCs and IONs inputs

Our main aim in this study was to characterize the two input streams to the cerebellum and compare their activity, in order to understand how they contribute to PC responses. In order to make some headway in this direction, we first asked to what extent, PC activity could be expressed as a weighted linear sum of the various GC and ION profiles we found. We started from the average response profiles of GC and ION clusters during the “steps” protocol to build a panel of regressors that could be used to reconstruct PC activity (Figure 7A). Then, we used multivariate linear regression with cross-validation and LASSO regularization to reconstruct the individual stimulus responses of each individual PC as a weighted sum of the GC and ION regressors plus an offset term (Figure 7A and Methods). As both these populations contact PCs with excitatory synapses, we constrained the model to have only positive weights. The regularization parameter was estimated using cross-validation and it provided sparsity in the weight matrix, ensuring that only regressors important for describing the PC activity got a non-zero coefficient. The details of the model and the fitting are described in Methods. The activity of most cells was successfully captured by our simple linear model, with around 57% of cells with a better-than-chance fit cost (Figure S7A). As expected, there was an inverse relationship between fit cost and cell reliability (Figure S7B).

**Figure 7:**
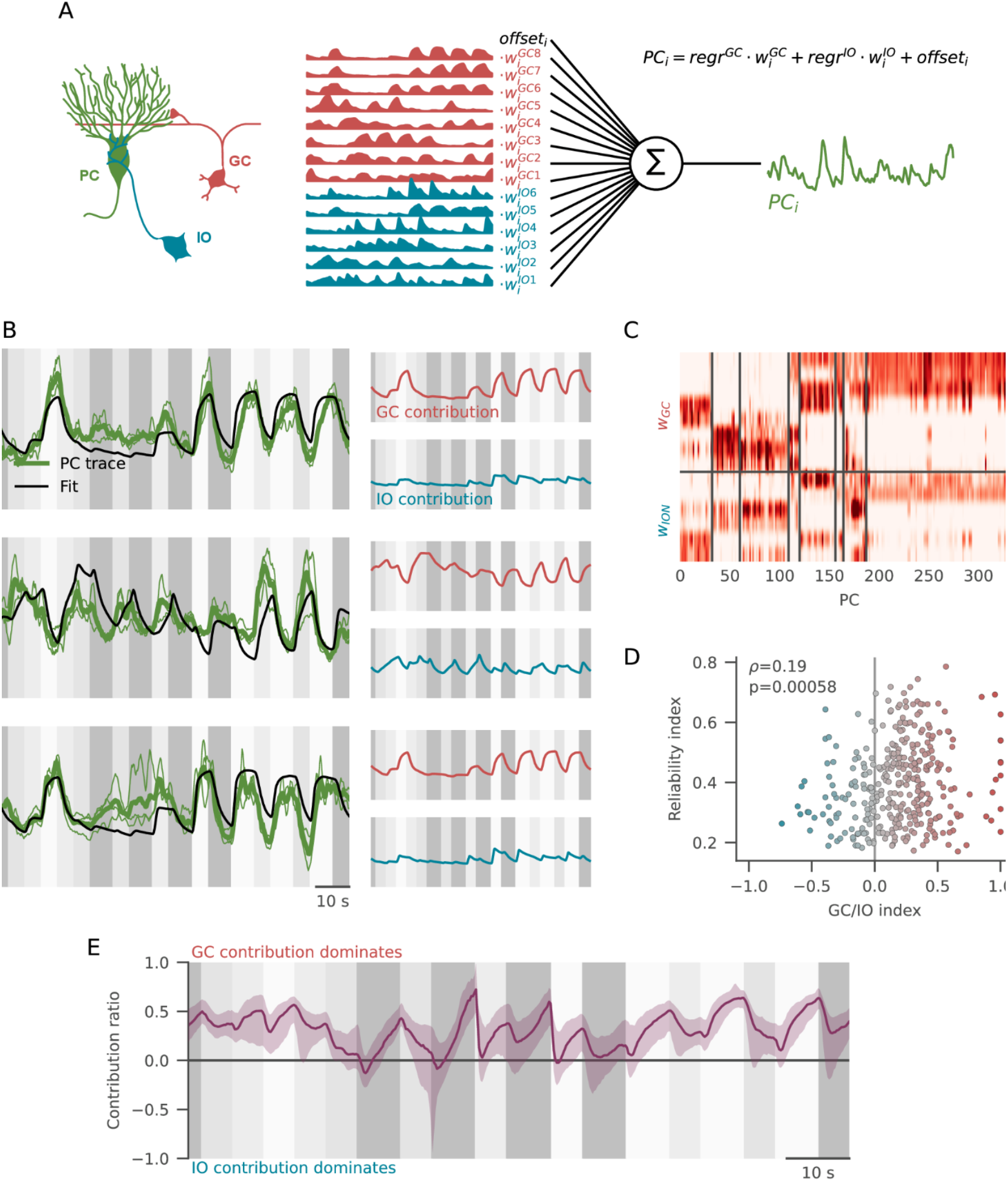
Reconstructing PC activity from IO and GC inputsA) Schema of the modeling approach. (Left) PCs receive inputs from IONs and GCs. Starting from GC and IO response clusters (center) we linearly combined them trying to reconstruct each PC activity (right). B) Examples of individual fits. (Left) Average PC response (thick green line) calculated on the individual test repetitions (thin green lines), and reconstructed trace from the model((black line). (Right) Red, trace reconstructed with GC coefficients only; blue: trace reconstructed with IO coefficients only. C) Matrix of weights assigned to each regressor (rows) for all PC ROIs (columns). D) Scatter plot showing the relation between the GC/IO weights ratio vs. cell reliability. Each dot represents the two values for a single PC cell, color coded by the GC/IO index. The response of PCs dominated by GC inputs are more reliable compared to cells dominated by IO inputs (Spearman rho: 0.19, p = 5.8*10-4). E) Evolution in time of the median contribution of either GC or IO clusters to the activity of PCs. Purple line: median contribution ratio trace across the entire PC population; purple shades: first-to-third quartile range.

The relative contribution of GC and ION regressors varied across the PC population. Figure 7B shows three examples of the fit obtained for three different PCs displaying different degrees of similarity with the GC or the ION responses. From the connectivity of the cerebellar circuitry, we would expect the number of non-zero weights for ION regressors to be lower than the number of GC regressors. This was indeed the case, although, surprisingly, also the number of required GC regressors required to predict PC activity was low (Figure S7C, 1.6 vs 2.8 average non-zero weights per PC for GC and ION regressors, respectively). The diversity in the distribution of the weights across PCs matches the clustering on the traces (Figure 7C) indicating that these functional classes identified in Figure 6 may arise from different connectivity of their inputs. To quantify the relative contribution of GC and ION clusters across the PC population, we computed for each cell a GC/ION ratio index going from −1 (only ION clusters with non-zero weights) to +1 (only GC clusters with non-zero weights, see Methods). The average GC/ION index was > 0 (number - error), with most cells having a stronger contribution from GC clusters compared to ION, confirming our qualitative observation that, when probed with our luminance stimulus, GC input dominates the activity profile of PCs.

Given the differences in the reliability indexes of the GC and the ION responses (Figure 2A-B), we would expect that PCs dominated by ION inputs would show a lower reliability when compared with neurons dominated by GC inputs. This was indeed the case: there was a significant positive correlation between the reliability index of the PC and the GC/IONs index (Spearman *ρ*=0.19, p=0.0006; Figure 7D). We had previously observed that whereas GC responses could be sustained in time, ION activity was more transient in nature, concentrated around stimulus transitions. Indeed, by computing the instantaneous contribution to PC activity from their GC and ION inputs across a trial, we observe that GC input dominates except during stimulus-OFF transients (Figure 7E).

In conclusion, we observed that PC responses can indeed be reconstructed as a linear combination of GC and ION responses profiles. Although the activity of most PCs was better predicted using loads on GCs regressors, the PC population exhibited a broad diversity of the GC vs ION loads, ranging from ION inputs only, to GC inputs only.

## Discussion

Here, we undertook a large-scale investigation of the representations of a simple visual stimulus in the olivo-cerebellar system of larval zebrafish at the single-cell level. Previous reports from our lab have been exploring the responses of individual cell types in the cerebellar circuitry (Knogler et al., 2017, 2019; Harmon et al., 2020; Felix et al., 2021). Building on this previous work, we have systematically explored the responses of IONs and GCs to changes in luminance. Furthermore, we recorded activity in PCs, which receive both GC and ION input, in order to understand how they could combine these signals.

### Stimulus state and transition coding in GCs and IONs

We observed that a large fraction of GCs were recruited by our sensory stimulation, strengthening previous observations that point against the notion of sparse coding in GCs (Giovannucci et al., 2017; Knogler et al., 2017; Wagner et al., 2017). Responses clustered into few groups which, nevertheless, displayed a diversity of response properties, varying in their sign, threshold, and saturation points. Intriguingly, some GCs were not clearly luminance-ON or luminance OFF but were excited by intermediate luminance and inhibited by strong luminance (and *vice versa*). This activity profile could be easily generated by the interplay between an excitatory input coming via mossy fibers and recurrent inhibition provided by Golgi cells, combined with different threshold and saturation levels in GCs (Bratby et al., 2017). However, we cannot exclude that such responses are not directly inherited by pre-cerebellar inputs (Barker et al., 2017). The diversity observed in these response types would suggest that the representation of the sensory input in the GC layer is not only dense but also allows to reliably decode the presented luminance even from the responses of small subsets of GCs.

On the other side, a smaller fraction of IONs showed consistent responses during our stimulation. Importantly, the responses of IONs were much more phasic in nature compared to GC, with a single peak following light-to-dark or dark-to-light transitions (or, less often, both). This observation is in agreement with the idea that climbing fibers report “error” or “salience” signals, that can result in cerebellar learning by inducing plasticity on GC to PCs synapses which convey contextual evidence about the state of the environment. As a result, while decoding the presented luminance from IONs activity was worse than GC activity-based decoding, IONs traces allowed for a more reliable discrimination of the stimulus transition times.

Together, these observations confirm the idea that parallel fibers from GCs provide contextual evidence to the cerebellum, while IONs code for sudden changes in the environment, a central tenet of models of learning in the cerebellum.

### Temporal coding in GC responses

The cerebellum is involved in the acquisition and timing of motor sequences. This is exemplified in classic paradigms of delay conditioning, where the conditioned motor response has to be precisely delayed from the onset of the conditioning stimulus. This requires that even the simple conditioning stimuli used in paradigms such as eye-blink conditioning need to evoke GCs responses that vary in time, so that selective reinforcement of some parallel fibers synapses can modulate the PC firing rate at specific intervals from the onset of the stimulus. Several mechanisms have been proposed to underlie such temporal patterning (Medina and Mauk, 2000; Yamazaki and Tanaka, 2009). On one hand, in delay line models, the stimulus-elicited activity is delayed by a fixed amount of time by variable number of synaptic connections in the pre-cerebellar neurons that impinge onto each GC, making the activity of GCs sparse in time, with variable onsets (Moore et al., 1989). On the other hand, in oscillatory models, the responses of GCs are supposed to be oscillatory, with a different characteristic frequency in each GC, so that they can sum up to form a basis for arbitrarily delayed PC responses (Gluck et al., 1990). Finally, in spectral models, the temporal patterning arises as a combination of varied membrane time constants in GCs and in inhibitory companion Golgi cells, that generate unique time constants for the on and off phase of each GC (Bullock et al., 1994; Bratby et al., 2017).

Although some work has shown hints of temporal coding in GCs (Kennedy et al., 2014) this has never been investigated at the whole GC population scale. Previous reports from our lab have reported no evidence of temporal patterning in GCs (Knogler et al., 2017) but they analyzed temporal patterning in recordings from a very limited number of neurons. In our experiments, we could find no evidence for temporally sparse or oscillatory activity, but we do see a continuum spectrum of delay in the responses, from fast onset, fast offset GCs to GCs whose activity was suppressed for several seconds before starting to ramp up. Interestingly, this was happening over a much longer timescale than what usually has been investigated in cerebellar studies. We report GCs whose onset followed by up to 5-6 s the stimulus onset, and whose activity was still increasing after 20 s of stimulus presentation. IONs, on the other side, were carrying very little temporal information about the ongoing stimulus. A decoding approach confirmed that accurate prediction of time occurred since stimulus onset was possible from GC activity but not from ION activity. Our data suggests that GCs can indeed provide a base for temporal coding in the cerebellum, and strongly supports the spectral timing model for temporally specific cerebellar learning.

### PCs integration of GC and ION activity

Finally, we aimed at describing the activity of PCs as a linear combination of IONs and GCs inputs. It has already been shown that such a linear modeling on afferent inputs can provide a good description of the PCs activity (Chen et al., 2017; Tanaka et al., 2019). Indeed, wWe could describe most of the PCs responses as a linear weighted sum of responses from IONs and GC clusters. As IONs are more active at stimulus transitions, PC dynamics is more driven by ION activity at transition times. Perhaps surprisingly, there was some heterogeneity in the amount of contributions from ION and GC clusters between cells, with some PC cells mostly showing ION-like activity and a majority of cells that were described better by GC regressors, hence suggesting that GC and ION inputs related to the same sensory modality do not necessarily converge over the same PCs. An obvious caveat of our study is the focus on calcium imaging dynamics in PCs, as calcium transients might be produced by different biophysical dynamics for ION-elicited complex spikes and simple spikes. However, it was recently shown that in larval zebrafish both complex spikes and bursts of simple spikes could be detected using the PC:GCaMP6s line (Knogler et al., 2017, 2019), and we could indeed observe both GC-like and ION-like activity in PCs.

In conclusion, our work provides the first parallel characterization of population responses of GCs, IONs and PCs in the cerebellum. This approach gives new insights into how stimulus features and timing are differently represented in the two converging cerebellar pathways and are integrated at the level of PCs, and could be leveraged in the future to mechanistically address the involvement of the cerebellum in processing sensorimotor signals.

## Acknowledgements

The authors thank Daniil A. Markov for help in initial stages of the project and sharing of analysis code.

LP and RP were funded by the Deutsche Forschungsgemeinschaft (DFG, German Research Foundation) through grant PO 2105/2–1. This work was funded by the Deutsche Forschungsgemeinschaft (DFG, German Research Foundation) under Germany’s Excellence Strategy within the framework of the Munich Cluster for Systems Neurology (EXC 2145 SyNergy – ID 390857198).

## Author contributions

OP, LP, and RP conceived the project and designed the experiments. OP acquired the behavior and two photon imaging data. OP analyzed the behavior data, and LP and OP analyzed the imaging data, VŠ did all the decoding analyses. OP and LP generated the figures. OP, LP and RP wrote the manuscript with help from VŠ.

## Data and code availability

All data and code to reproduce the manuscript has been made public:

- All the source data can be found at https://doi.org/10.5281/zenodo.7071734
- Analysis code including code that generates all the figures in the paper can be found at https://doi.org/10.5281/zenodo.7071725

## Methods

### Zebrafish lines

The following experiments were performed with 6-8 days post-fertilization (dpf) zebrafish (*Danio rerio*) larvae. Before experimentation, larvae were kept on a 14h light/10h dark cycle and at a constant temperature of 28°C. Three types of transgenic lines in a nacre (*mitf*-/-) (Lister et al., 1999) genetic background were used for functional imaging experiments. For GCs and IONs imaging, a modified fast calcium indicator under the UAS promoter UAS:GCaMP6fEF05, see (Felix et al., 2021)) was expressed either under a GC-specific Tg(gSA2AzGFF152B) (Takeuchi et al., 2015)) or ION-specific (Tg(hspGFFDMC28C), (Takeuchi et al., 2015)) Gal4 promoter. For PC imaging, GCaMP6s was expressed under a direct PC promoter (Tg(PC:GCaMP6f), (Knogler et al., 2019). For behavioral experiments, the Tuepfel long-fin (TL) wild-type strain was used.

All experimental procedures were performed in accordance with the guidelines from the Regierung von Oberbayern, and following protocols approved by the Max Planck Society.

### Stimuli

All stimuli were presented to the lower part of the visual field of the fish, to cover the whole base of the dish where the larvae were embedded or swimming. These stimuli consisted of different combinations of sharp luminance changes, and their display was controlled with Stytra (Štih et al., 2019). For the experiments presented in this study, two different protocols were used. The “steps” protocol consisted on 5 s luminance steps of three different brightness levels (corresponding to 5%, 20% and 100% of the maximal luminance, chosen to map quasi-logarithmically the dynamic range of the projector) with 7 s inter-stimulus intervals. All the possible luminance transitions between the different levels were sampled in each repetition. The “flashes” protocol consisted of 3-, 7- and 21-s flashes of maximal brightness, interspaced by 7 s inter-stimulus intervals. The projectors were calibrated before the design of the stimuli: the lowest and highest luminances (corresponding to 0 and 255 pixel brightness values) were 0.3 lm and 55.7 lm respectively.

### Freely swimming behavioral experiments

For behavioral experiments, 6-7 dpf fish larvae were placed 3 at the time in about 1 cm of water in a rectangular-shaped arena cut in a 1% agarose matrix in 88 mm Petri dishes. The dish was placed on top of a light-diffusing screen mounted on a clear acrylic support, illuminated from below using an array of IR LEDs (setup fully described in (Štih et al., 2019) and in Stytra documentation). Larvae were tracked online at 500Hz using a high-speed camera (Ximea MQ013MG-ON) with Stytra. Visual stimulation was displayed from below using an Asus P2E microprojector and a cold mirror (Edmund Optics). A red long-pass filter (Kodak Wratten No. 25) was placed after the projector to match the absolute luminance values and wavelength of the imaging experiments stimulation. In freely swimming experiments we used the “steps” protocol, presented a total of 36 times (1 h total duration).

### Functional imaging experiments

For functional imaging, larvae were placed in 35 mm Petri dishes and embedded in 2.5% agarose. The dishes were placed onto an acrylic support, on top of a light-diffusing screen, and either the cerebellum or the inferior olive were systematically imaged with a custom-built two-photon microscope. For excitation, a TiSapphire laser (Spectra Physics Mai Tai) tuned to a 905 nm wavelength was used. Visual stimuli were projected from below at a rate of 60 frames per second using an Asus P2E microprojector, and a red long-pass filter (Kodak Wratten No. 25) was placed in front of the projector. Imaging frames were acquired every 248.88 ms (for cerebellum imaging) or 246.38 ms (for inferior olive imaging). The protocol was presented 6 times while acquiring every plane, then the focus was shifted ventrally 1.5 μm (for cerebellum imaging) or 1 μm (for inferior olive imaging) and the process was repeated. For every fish about 50-70 planes were imaged.

### Image processing

Alignment. 2P data were first aligned with a plane-wise rigid transformation. A reference image for each plane was computed as the average of n frames, and displacement of each individual plane from the reference was found through image cross-correlation and corrected with rigid translation. Frames with a correlation peak > 10 microns from the center, usually coming from motion artefacts, where set to nan and discard in following analyses. To correct for shifts happening between planes, and similar procedure was followed calculating shifts from one plane to the next using cross correlation between consecutive planes averages.

ROI extraction: For ROI extraction, we used a combination of approaches. For GCs and PCs, as the number of cells is high, manual segmentation would be unfeasible; but, as responsive cells were generally reliable, it was possible to reconstruct individual ROIs spanning multiple planes thanks to the similarity of the signal between planes. For IONs, the sparse activity made it difficult to match ROI activity across planes based on correlation, but the low number of cells (∼80 per fish) makes manual segmentation feasible.

For automatic ROI extraction, we used exactly the same iterative procedure described in (Markov et al). Briefly, a spatial correlation map was computed where each pixel was assigned a value corresponding to the correlations over time between its fluorescence and average fluorescence of the 8 neighbor pixels. Then, starting from the yet unassigned pixel with the highest correlation value (seed), neighbor pixels were included in the ROIs if their fluorescence correlation with the average fluorescence of the ROI grown so far passed a certain threshold.

For manual ROI extraction, a custom python GUI was used to draw ROIs on the correlation maps.

### Response analysis

All the analysis on the obtained traces was done in Python; Jupyter notebooks generating all the figures of the paper are available in the linked repository.

#### Reliability index and filtering

to find a measure of responsiveness to the presented stimulus as independent as possible from the specific response profile, we calculated for each ROI we calculated the average correlation of the responses across all individual presentations of the stimulus. The distribution of the obtained correlations had a peak close to zero and a second peak (or a long tail) of positive correlations corresponding to responsive cells. To use an objective criterion to select responsive cells, we used Otsu’s method from the scipy package to set a threshold on the obtained histogram.

#### Hierarchical clustering

Next, we calculated the mean response for each responsive ROI and we used the ward algorithm for hierarchical clustering (from the scipy package) to cluster them. We manually specify a cut on the clustering tree to obtain the discrete clusters that we use in the rest of the analysis. We arbitrary decided for a threshold that was low enough to include all clusters with qualitatively different changes in their response properties

#### Regressor analysis

For regressor-based analysis, we manually designed a set of regressors starting from either luminance profiles with a gamma correction of 1, ½ or 2; the difference between the luminance profile with gamma 2 and with gamma 1 (imitating intermediate luminance responses); and on, off, and combined on and off transitions. The obtained regressors were then convolved with a kernel of (n) s, to match the temporal response of GCaMP6fe05. For each cell, Pearson correlation with all regressor was computed.

#### COM sorting

The COM was defined as the point in time where the integral of the neuronal trace reached half of its total integral value. To cross-validate the plots, its value for each individual ROI was estimated based on the half of neuronal responses that were not used for the plotting (Figure S5A).

#### Decoding

To decode different information from the cell activities, we used standard methods from the Scikit-Learn Python package (Pedregosa et al., 2011). Every decoding analysis was trained on 10 randomly chosen trials and tested on two others (only cells with data from 12 or more trials were used in the analysis). Ridge regression was used to decode the time since stimulus onset (Figure 5E). Kernel-based support vector regression (SVR) was used in Figure 3D. Grid search over the one regularization parameter was performed in all cases by leaving out each of the 10 training trials. For all decoding analyses we have to caution that the population activity used to decode the populations is constructed. Neurons were sampled from different repetitions and animals, therefore destroying correlations between neurons. In many cases, such correlations, often termed noise correlations (because these remain unaccounted for after taking out stimulus responses) can have a significant impact on the decoding quality (Moreno-Bote et al., 2014). The decoding analysis is provided together with the rest of the analysis code.

#### Model fitting

For the PC modeling analysis, we used the data acquired with the “steps” protocol. First, we calculated average responses from all the GCs and IONs clusters (8 clusters of GCs, and 6 clusters of IONs). To address potential differences in time constants between calcium sensors, we deconvolved average responses using the GCaMP6fe05 kernel, and convolved using the GCaMP6s kernel to match the sensor used in PCs. The resulting average traces were normalized to be strictly positive and with integral 1. Then, the traces from luminance responsive PCs were Z-scored on a trial-by-trial base, high-pass filtered with a very low cutoff frequency (1/80 Hz) to remove slow drifts, and smoothed with a 3 pts mean boxcar window to reduce noise.

The function that was optimized for each cell was the following:

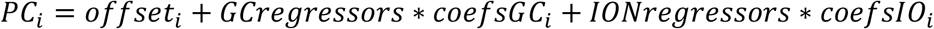

Where:

- *PC*_*i*_ is the trace for the i^th^ PC cell;
- *GCregressors,IONregressors* : are the matrix of GC and ION regressors

The optimization was done on the following parameters:

- *offset*: a constant term, bound to be between −5 and 5
- *coefsGC*,: coefficients for GC regressors, bound to be between 0 and 1000
- *coefsION*: coefficients for ION regressors, bound to be between 0 and 1000

The large coefficient boundaries come from the different normalizations applied on regressors - norm - and on trace −Z scoring.

For each cell, 2 stimulus repetitions (“test” data) were left out to be the final test traces for the analysis and excluded from the entire fitting process. The remaining repetitions (4 or more, for cells spanning multiple planes) were used to find the regularization term and doing the fit.

To find the L1 regularization lambda parameter, we used leave-one-out cross-validation to calculate fit costs over a logarithmic sweep of lambdas between 10^-7 and 10^-2. We trained the model on all the fit traces but one, we calculated the fit cost on the remaining trace and took the average value across all the left-out traces. Then, we calculated the average lambda and used it for all cells, to allow for a more appropriate comparison of the fit parameters across cells.

After the cross-validation of regularization lambda, the parameters were fit over all the “fit” repetitions. The “test” repetitions were then used to estimate final fit performance and for all the plots in Figure 7.

### Behavioral analysis

To analyze the behavioral responses to the luminance transitions, we started from raw velocity traces saved by Stytra. As zebrafish larvae swim in discrete events called bouts, the fish speed was thresholded to extract individual bout events. The time occurrence of each bout event relative to the presented stimulus was binned, a histogram of bout times was obtained for each fish (Figure S1A). To assess whether the decrease of bout probability after each luminance off transition was statistically significant, we bootstrapped a distribution of bout probabilities from the histogram of each fish, and compared the average bout probability 1.5 s after the off transition with the 1st percentile of the bootstrapped distribution (Figure S2C). In all but three fish, the post-transition probability was lower than the 1st percentile of the histogram (Figure S2C-D).

## Supplementary Figures

**Figure S1.**
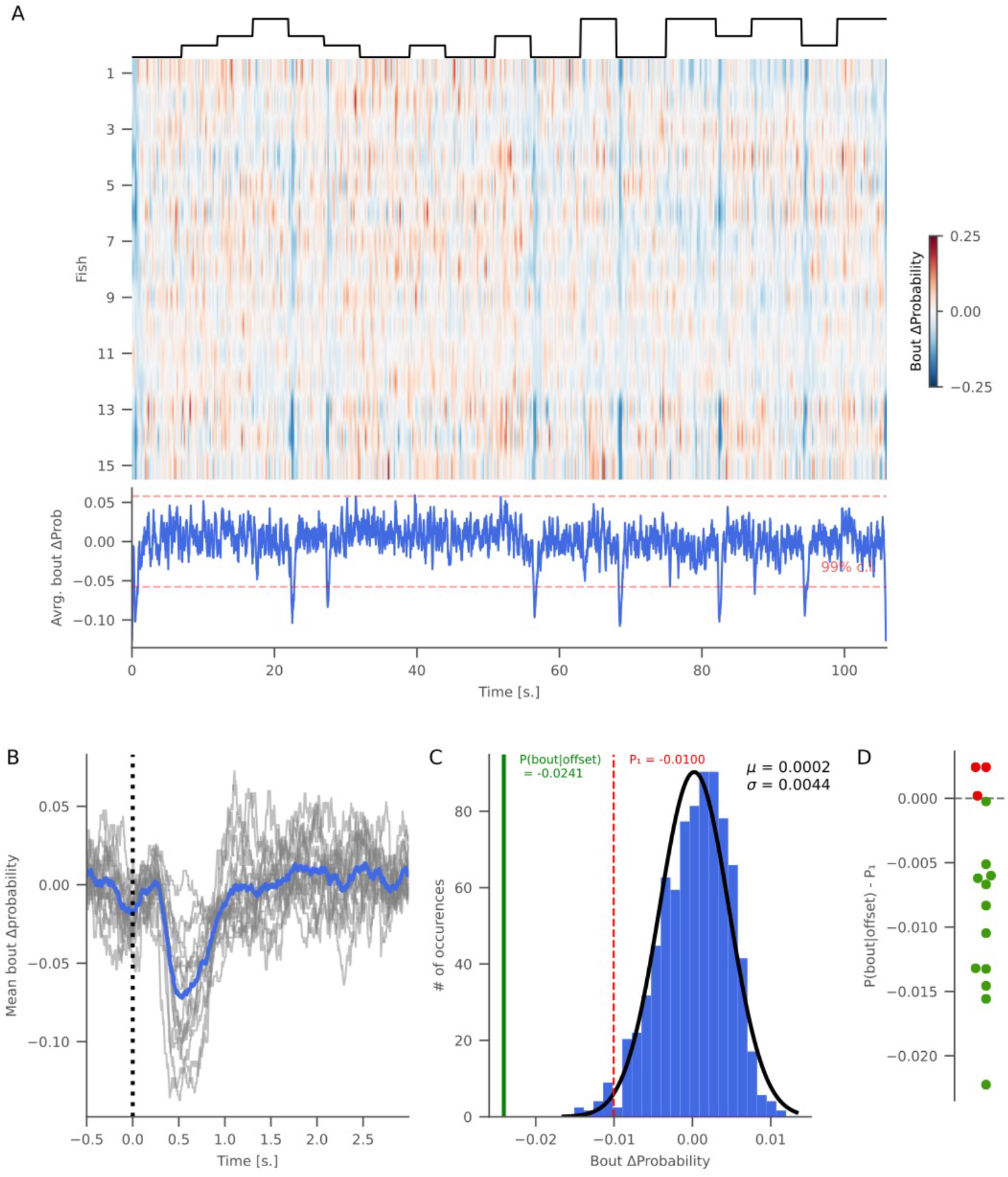
A) Average normalized bout probability for each individual fish across all stimulus presentations (heatmap), and average probability across all fish (blue line). B) Bout probability changes on a 3.5 s window around luminance offsets. Average bout probability across all off transitions is shown for each fish in gray lines, and the average bout probability across all fish is plotted in blue. C) The average bout probability at offset (during the 1.5 s following luminance changes, green line) across all fish, compared to the 1st percentile (red line) of a dataset generated via bootstrapping. D) Difference between the bout probability at offset and the 1st percentile for the bootstrapping analysis performed individually on each fish. Fish labeled in green correspond to animals where the bout probability was smaller than the 1st percentile.

**Figure S2.**
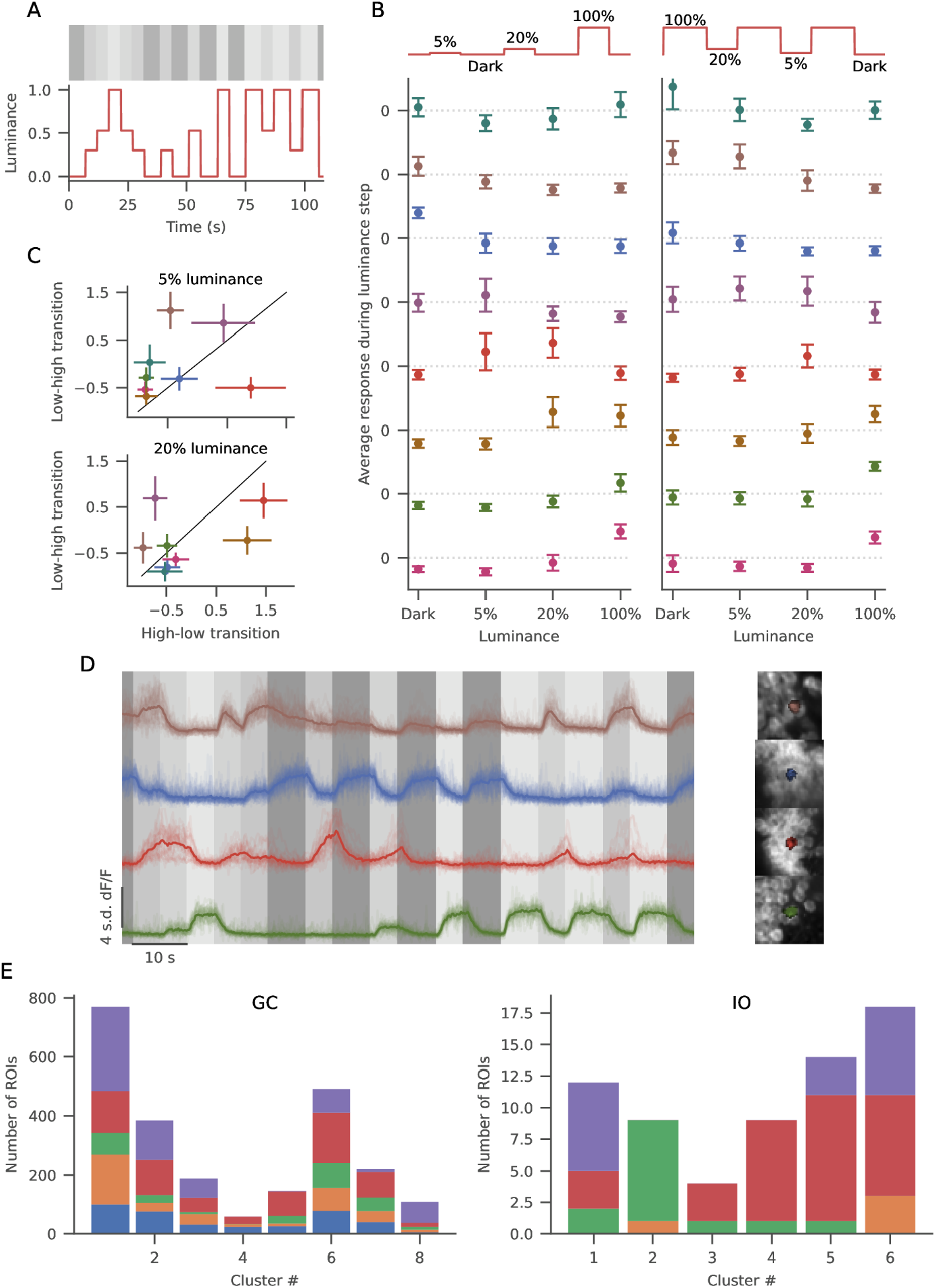
A) Schema of the stimulus presented, with the color scheme used for all figures mapped on top of its luminance profile. B) Average fluorescence during the upward luminance steps from minimum luminance (left) and during downward luminance steps from maximum luminance (right) for each GC cluster. C) History dependence of luminance responses for all GC clusters. Average normalized fluorescence during the presentation of the same two intermediate levels of luminance (low intermediate: above, high intermediate: below), compared in epochs when it was reached from a higher (x values) or lower (y values) luminance level. Clusters that deviate from the diagonal are the ones showing the strongest temporal history dependence (color-coded according to Figure 2A).D) Example of GC responses from various clusters, with individual stimulus repetitions (thin lines), average (thick line), and the ROI morphology (on the right). E) Contribution of individual fish to the observed clusters for GCs and IONs. Each color corresponds to one fish.

**Figure S3.**
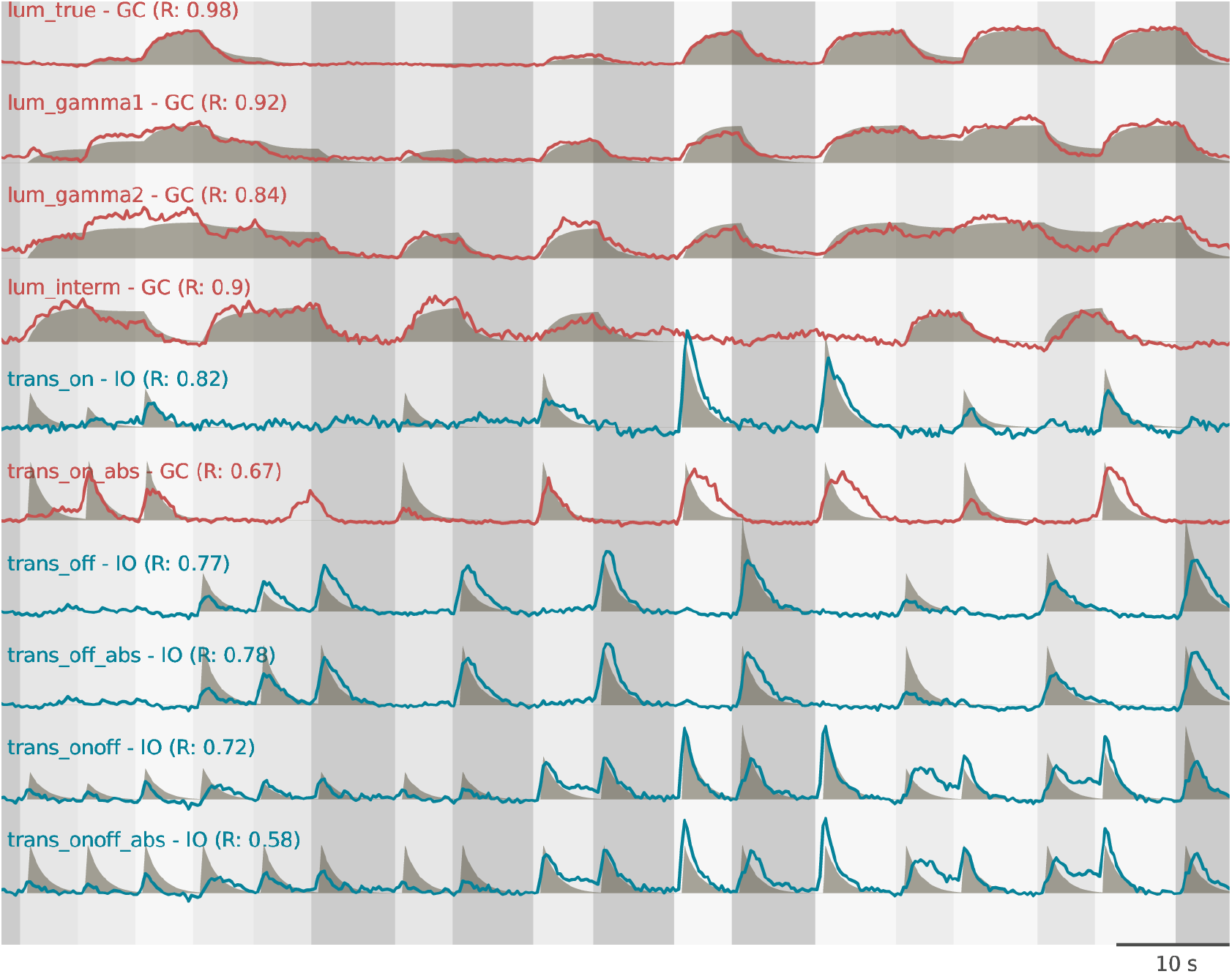
All regressors used in the regression analysis (shades), and the best scoring ROI for each regressor (lines). While for luminance-related regressors the highest correlation values were always from GCs, for transition-related regressors most of the best scoring ROIs were from IONs.

**Figure S4.**
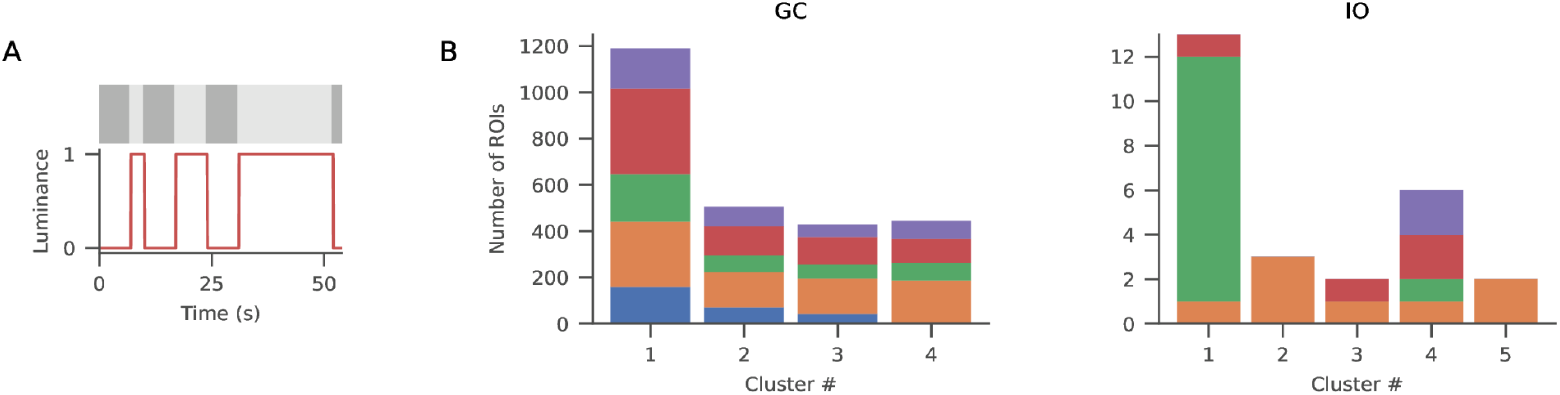
A) Schema of the stimulus presented, with the color scheme used for all figures mapped on top of its luminance profile. E) Contribution of individual fish to the observed clusters for GCs and IONs. Each color corresponds to one fish.

**Figure S5.**
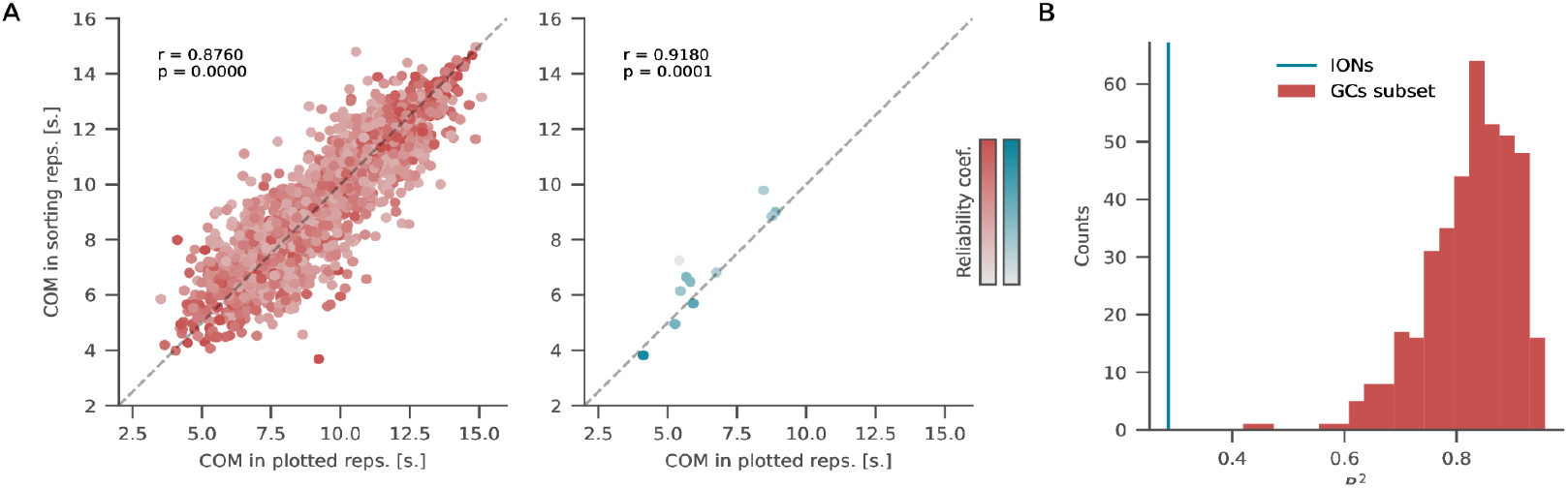
A) In order to cross-validate our sorting of ROI responses, the COM for each neuron was calculated based on the average response during half of its repetitions, and the other half of the repetitions were used to plot the figures shown in Figure 5. Figure S5A shows, for each ROI, the COM calculated on each half of the repetition. B) Histogram of the R2 values between predicted and actual time from stimulus onset, as decoded from 200 subsamples of GCs. The blue line marks the R2 value obtained using the same number of IONs.

**Figure S6.**
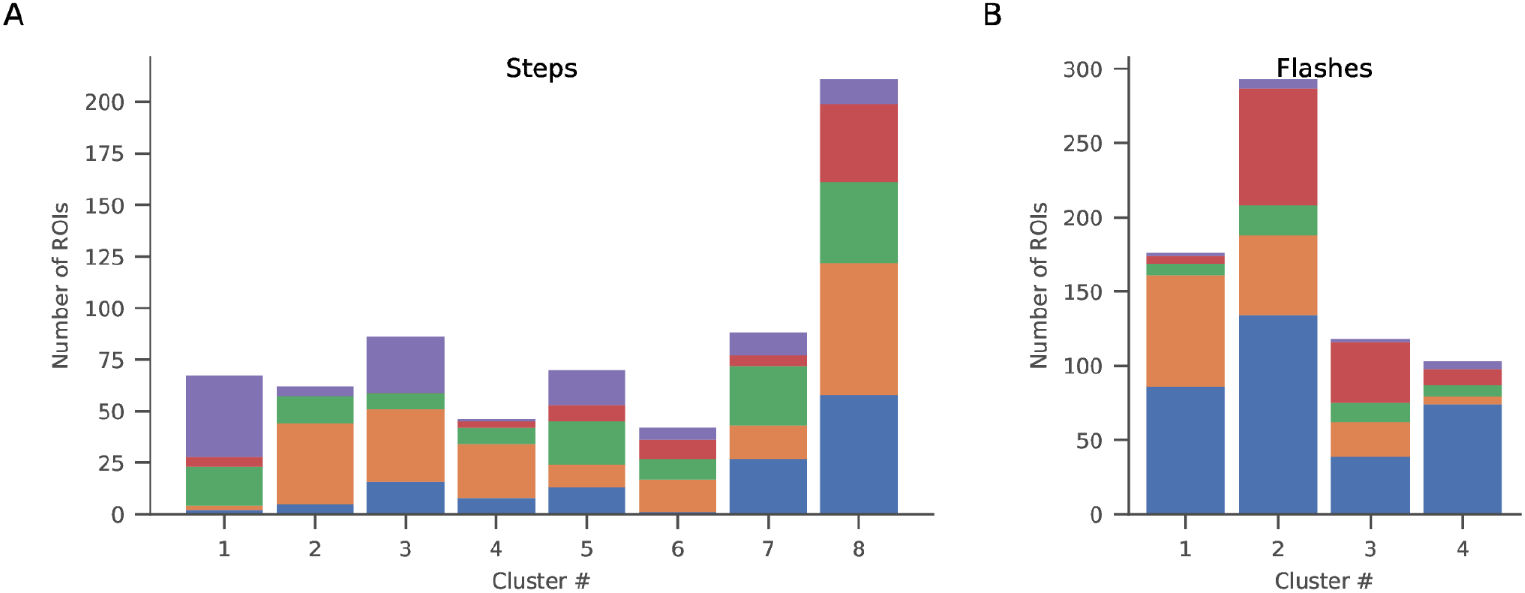
A) Contribution of individual fish to the observed clusters for PCs in the steps protocol. Each color corresponds to one fish. B) Contribution of individual fish to the observed clusters for PCs in the flashes protocol.

**Figure S7.**
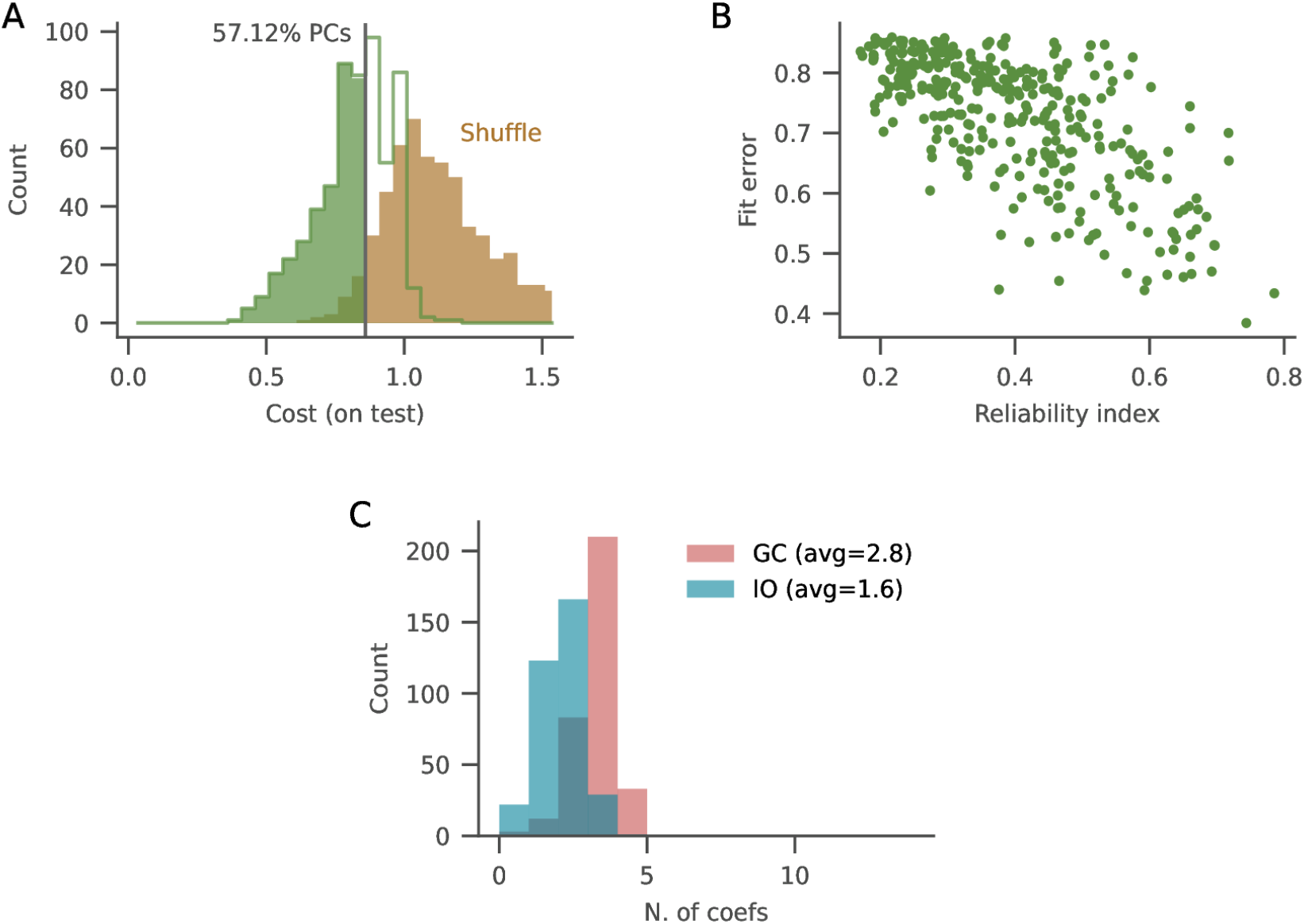
A) Histogram of costs (L^2^) on the test traces from all PCs (green line), compared to a shuffle distribution (brown shade) that was obtained calculating costs after a random regressor-wise reshuffling of the weight matrix (i.e., all values from each individual regressor were kept and reshuffled in new random combinations). A threshold was defined to ensure that only 5% of these random fits could have a lower cost. The green shade indicates the data that were kept after such selection. B) Correlation between the reliability index and the fit error, including only PCs for which the fit was considered better-than-random. C) Distribution of the number of non-zero weights for IONs (average: 1.6 weights) and GCs (average: 2.8 weights) regressors.

